# A Preliminary Application of Bird Surveys Using Environmental DNA Technology: in Futian Mangrove Nature Reserve

**DOI:** 10.1101/2025.01.13.632664

**Authors:** Song Zhang, Xiao Ye, Jiang Chen, Xiao Jun Liu, Tao Jin

## Abstract

Futian Mangrove Nature Reserve is a vital habitat for birds in Shenzhen city of China, and the diversity of bird species is of considerable concern. Traditional surveys depend on direct observation and expert assessment, necessitating significant manpower for tracking. This study utilized environmental DNA metabarcoding technology to gather 128 environmental samples of water, soil, and air, along with samples of bird feces and feathers collected during the summer and winter of 2023. A total of 72 bird operational taxonomic units (OTUs) were identified. The results revealed variations in bird detection across different environmental samples, with air samples yielding 52 bird OTUs, water samples 29, and soil samples 23. Detected bird OTUs were notably higher in winter samples compared to summer. Additionally, the average number of bird OTUs detected in air samples during summer and winter was 14.9 and 10.1, respectively, significantly exceeding those found in the corresponding water and soil samples. The identification of bird feces and feather samples confirmed the presence of the identified bird species in the environmental samples. This study concludes that environmental DNA technology can serve as a valuable complementary method for monitoring bird diversity, with different environmental samples showing distinct preferences in bird detection.

## 1 Introduction

Environmental DNA (eDNA) technology leverages DNA released into the environment, rather than relying on the direct capture of organisms(1,2). This method typically involves collecting environmental samples—such as water, soil, and air—enabling efficient and non-invasive biodiversity surveys(3). In recent years, as eDNA technology has matured, it has been increasingly applied in biodiversity assessments across various animal groups, including fish, insects, and mammals, with a particular focus on aquatic species(4,5). By extracting DNA from these environmental samples and amplifying gene fragments using universal primers that target specific mitochondrial genes (such as 12S rRNA, 16S rRNA, and COI), researchers can generate high-throughput sequencing data. The sequences undergo standardized operational taxonomic unit (OTU) clustering, and then the unique OTUs are compared to established databases, ultimately revealing the taxa present in the samples. Compared to traditional biodiversity survey methods that rely on morphological identification, eDNA technology offers several advantages, including ease of sampling, environmental friendliness, and high precision and sensitivity(6).

Birds have always been a significant focus in biodiversity surveys due to their high diversity. In China, approximately 1,400 bird species have been recorded, many of which are migratory. Historically, bird surveys have been conducted primarily by professional research teams, with contributions from birdwatching enthusiasts serving as valuable supplementary data. Such surveys heavily depend on the observers’ expertise, as many bird species are small and can only be observed from a considerable distance, particularly waterbirds. Several years ago, researchers began exploring the use of eDNA technology for bird monitoring, demonstrating its feasibility(7,8). However, compared to its application in fish and other groups, eDNA utilization in bird surveys remains in the exploratory phase, with critical operational procedures—such as sample collection and primer selection—requiring further optimization and standardization.

Futian Mangrove Nature Reserve, located on the northeastern shore of Shenzhen Bay, features well-developed natural mangroves. This reserve serves as a vital habitat and feeding ground for many wintering birds, including the endangered black-faced spoonbill, and is also a breeding ground for various local species. In this study, we collected three types of environmental samples—water, soil, and air— along with small quantities of feces and feathers within the reserve, utilizing eDNA technology to investigate bird diversity. The objectives of this study are to assess the feasibility of eDNA technology for bird surveys within the reserve.

## 2 Materials and Methods

### 2.1 Overview of the Study Area and Sampling Locations

Futian Mangrove Nature Reserve, located in southern Shenzhen (114°03′E, 22°32′N), is the smallest national nature reserve in China, covering 367.64 hectares, which includes 139.92 hectares of land and 227.72 hectares of intertidal zones. The reserve enjoys a mild climate, with an average annual temperature of approximately 22°C and average annual precipitation of 1927 mm. Classified as having a South Asian subtropical marine monsoon climate, it lies along the migratory route of Siberian-Australian migratory birds. Each year, tens of thousands of migratory birds stop here, making it a critical habitat and waystation for species in the eastern hemisphere(9). The mangrove ecosystem comprises three main subsystems: intertidal zones, mangroves, and aquaculture ponds, collectively providing vital habitats and foraging areas for birds(10).

Sampling sites were strategically selected in areas of frequent avian activity, including 10 semi-saline ponds, named as Pond 1 to Pond 10, one freshwater pond, and three intertidal areas (the birdwatching pavilion, Fengtang River estuary, and Shazui wharf), resulting in a total of 14 sampling points, as shown in Figure 1. The samples collected included environmental samples (water, soil, air) and biological samples (feces and feathers), serving as the DNA sources for bird barcoding identification.

**Figure 1:**
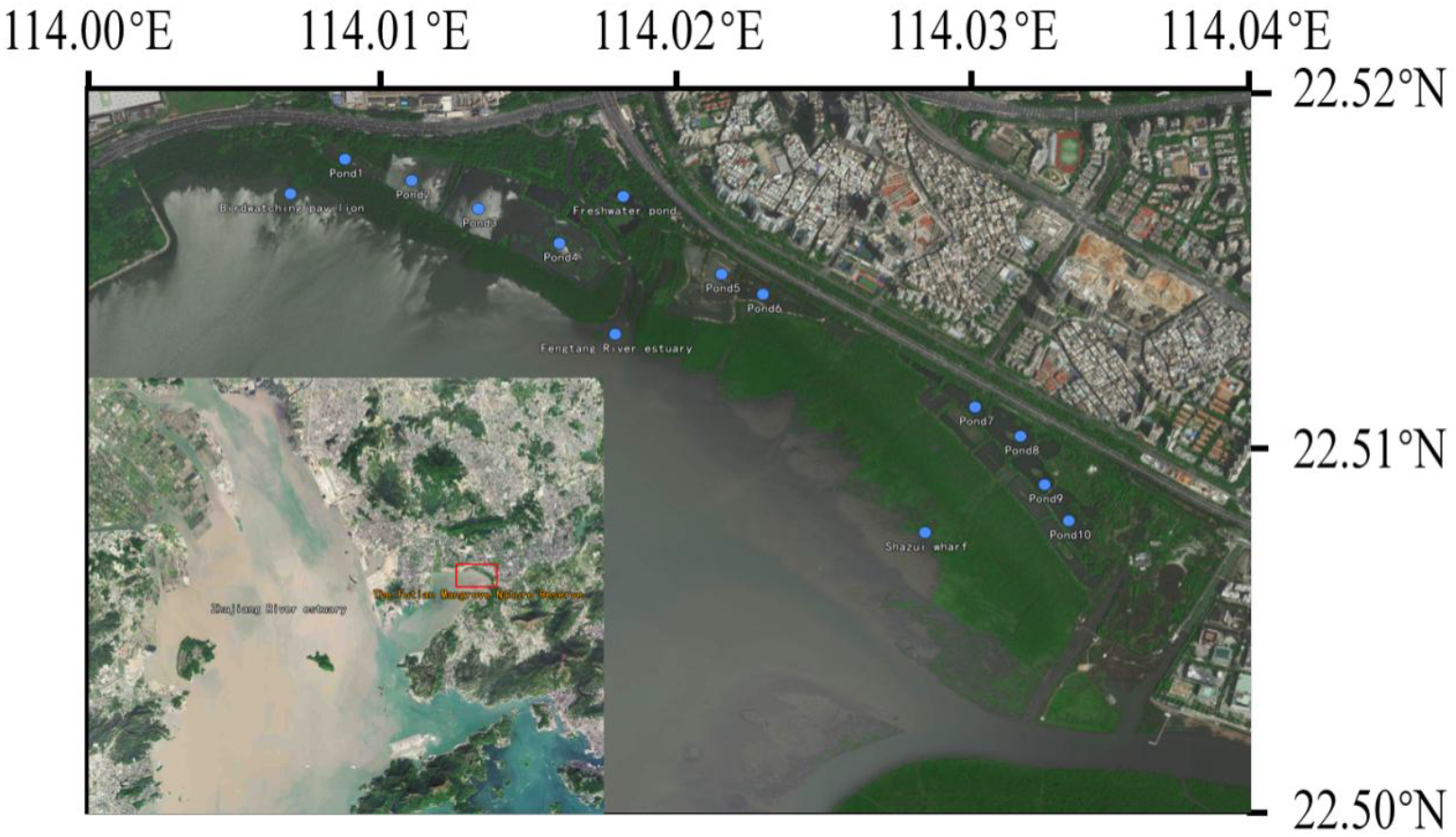
Distribution map of sampling sites.

### 2.2 Sample Collection

Samples were collected in summer (April to July 2023) and winter (November 2023 to January 2024), with a total of 128 samples collected primarily from the 14 designated sites. In summer, 21 water samples, 22 soil samples, and 20 air samples were collected. In winter, 22 water samples, 23 soil samples, and 20 air samples were obtained, with all sample types evenly distributed across the 14 points. Bird biological samples were collected randomly, primarily in winter, including 18 fecal samples and 13 feather samples. To avoid identifying birds from a single source, distinct morphological and color differences among the samples were noted.

Water samples were collected using a sampler in areas with a high concentration of water birds, ensuring significant distances between collection points at the same site. Samples were collected in 1L bottles, which were rinsed with 95% alcohol prior to use and then rinsed twice with site water. After sealing, samples were stored in an ice bath and vacuum-filtered within 8 hours using a 0.45 μm cellulose acetate filter (CA, Duohong), then stored at -20°C for future DNA extraction.

Soil sampling points were selected in areas of high bird activity, either on the shore or under trees. Soil was collected using a five-point sampling method, where samples from different locations at a point were combined. The collected soil was ground into powder using a soil crusher, mixed on a sterile cloth, and stored in sample bags at -20°C for future DNA extraction.

Air samples were collected using a mid-flow particulate matter sampler (JH-2010 model), positioned 1 meter above the ground. A 90 mm, 0.45 μm glass fiber filter was utilized, with an airflow rate set to 120 L/min for 4 hours. Different sampling points were at least 300 meters apart. The membranes were stored in centrifuge tubes at -20°C for future DNA extraction.

Bird fecal samples were randomly collected within the reserve and placed in centrifuge tubes, preferably selecting single droppings found on leaves or pathways. Feces from the intertidal zone or shore were collected from the upper portion to avoid contamination. Samples were categorized into single droppings (one sample for individual testing) and mixed samples (combined from several single droppings for pooled testing). Feather samples were selected to be relatively fresh and containing follicles, with each feather individually bagged.

### 2.3 DNA Extraction, PCR Amplification, and Sequencing

DNA from water and air filters was extracted using the TIANamp Marine Animals DNA Kit, while the ALFA-SEQ Advanced Soil DNA Kit (mCHIP) was used for soil, fecal, and feather samples. Universal primers designed by Wang et al. were employed to amplify mitochondrial 12S rRNA gene fragments(11). The primer sequences are as follows: V12S-U-F: 5’-GTGCCAGCNRCCGCGGTYANAC-3’ and V12S-U-R: 5’-ATAGTRGGGTATCTAATCCYAGT-3’. PCR amplification was conducted in a total reaction volume of 50 μL, which included 25 μL of Mix premix, 2 μL of each forward and reverse primer, 30 ng of DNA template, and nuclease-free water to fill. The amplification program consisted of an initial denaturation at 95°C for 10 minutes, followed by 36 cycles of denaturation at 95°C for 30 seconds, annealing at 56°C for 30 seconds, and extension at 72°C for 30 seconds, concluding with a final extension at 72°C for 5 minutes. After extraction, purification, and quantification of PCR products, sequencing libraries were prepared with standard adapters on the Illumina Nova 6000 platform to obtain DNA sequence data.

### 2.4 Data Analysis

Sequencing data processing began with the quality control of FASTQ files generated from the amplification of each sample using the Fastp tool (v0.12.4)(12). Sequences with an average base quality score below 20, lengths shorter than 100, and those with more than 3% ambiguous bases were removed. The quality-controlled paired-end FASTQ data were then assembled using USEARCH software (v11.0.677), requiring overlapping regions of 16 bases without mismatches(13). Subsequent quality control of the assembled sequences was performed with VSEARCH(14) software, where primer sequences at both ends were removed, retaining sequences between 152 and 182 bp. All assembled sequences from all samples were merged, and USEARCH was used for dereplication (fastx_uniques), chimera removal (uchime3_denovo), and OTU (Operational Taxonomic Unit) calculation. Sequence counting for each sample across all OTUs was performed using the usearch_global module, resulting in an OTU table for all samples. To account for sequencing errors, cross-contamination, and algorithmic biases, OTUs with relative abundances below 0.05% were set to zero(15,16).

### 2.5 Taxonomic Annotation

The resulting OTU sequences were subjected to local alignment (BLAST) against the NCBI database (https://www.ncbi.nlm.nih.gov/) and the BOLD database (https://www.boldsystems.org/) to retrieve taxonomic information of the best-matching reference sequences for each OTU. OTUs with similarity greater than or equal to 99% were assigned species-level annotations, those with similarity between 95% and 99% were assigned genus-level annotations, and OTUs with similarity below 95% were discarded. Additionally, the annotated bird species were cross-referenced with the “2023 Edition of the Shenzhen Bird List” published by the Shenzhen Birdwatching Association to ensure the accuracy of the detection results.

## 3 Results and Discussion

### 3.1 Analysis of Birds in Environmental Samples

Sequencing data from the 128 collected environmental samples revealed a total of 72 bird operational taxonomic units (OTUs)(Figure 2). Among these, 48 OTUs were identified at the species level, as detailed in Table 1, while the remaining 24 were classified only at the genus level, mainly due to a lack of reference sequences for certain bird mitochondrial 12S rRNA gene. The identified species comprised 38 forest birds and 34 water birds, indicating a balanced distribution between the two groups.

**Table 1:**
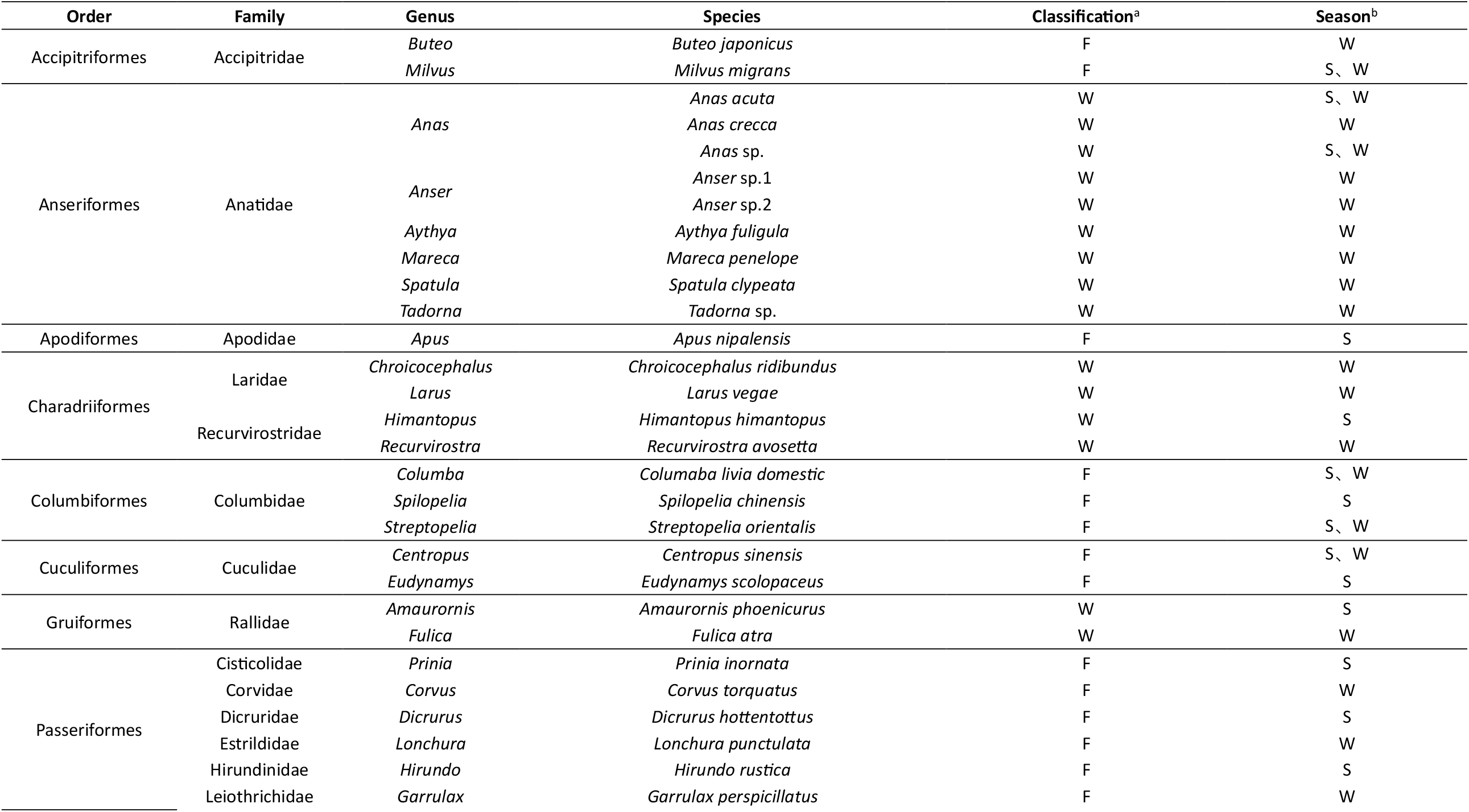
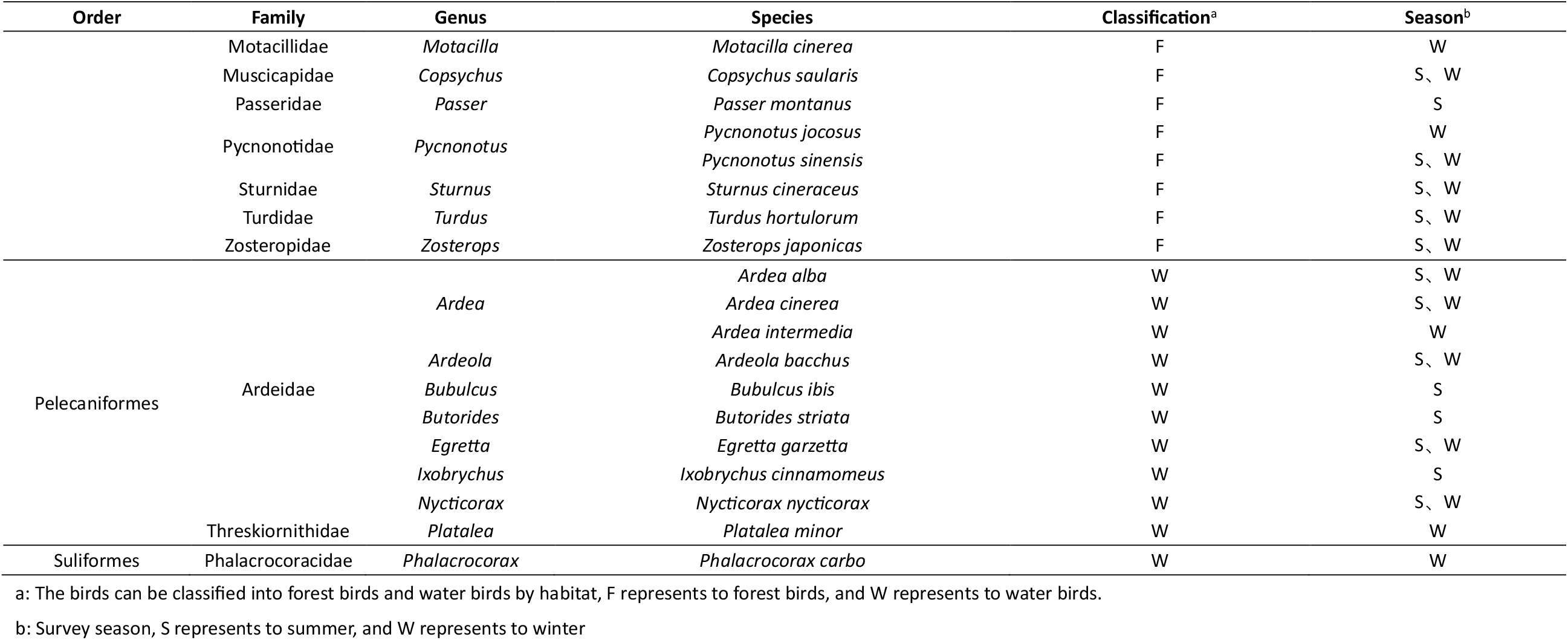
Bird species detected by eDNA in Futian Mangrove Nature Reserve in 2023.

**Figure 2:**
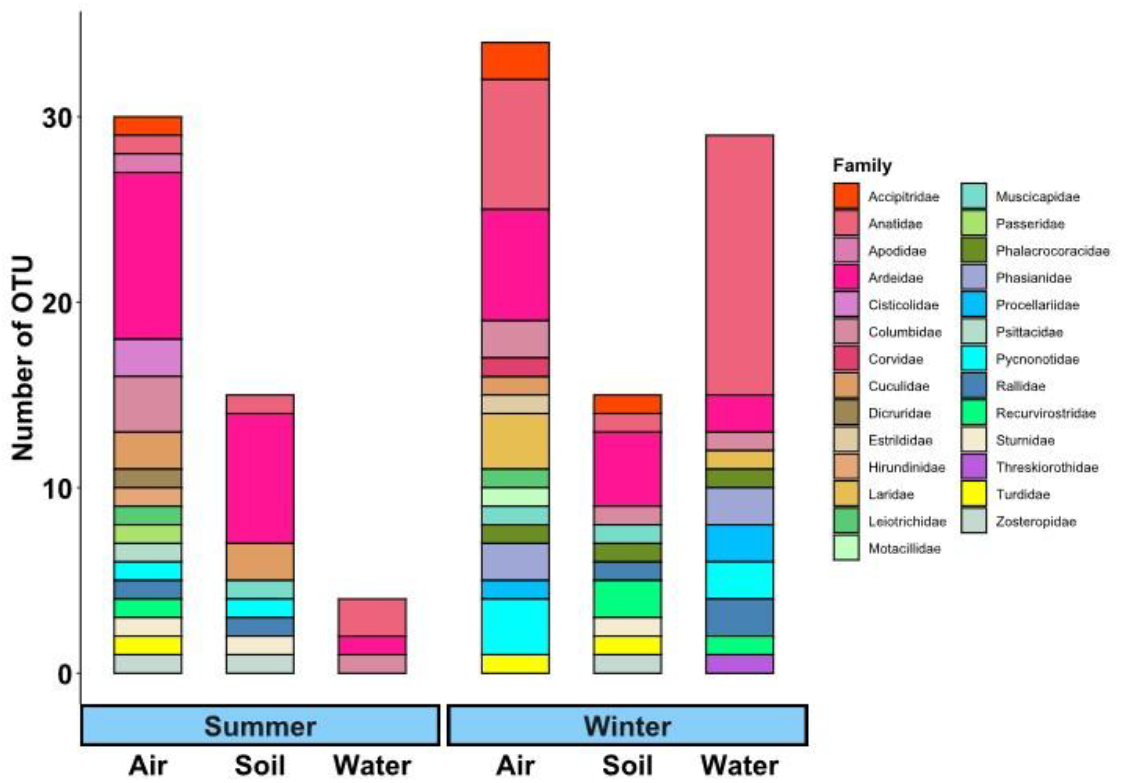
Bar chart of bird species composition detected in three types of samples during summer and winter. Different bars represent the number of bird species at the family level detected in samples from different seasons, with different colors indicating different families.

Further analysis of the environmental samples highlighted that air samples yielded the highest species diversity, detecting 52 species (28 forest birds and 24 water birds). 29 species were identified from water samples, consisting of 18 water birds and11 forest birds, reflecting the preference of water birds for aquatic environments. Soil samples recorded the fewest species, with 23 species(11 forest and 12 water birds). Some species were detected across multiple sample types, as illustrated in the Venn diagram in Figure 3B. Notably, the winter season resulted in a higher detection count of 51 species compared to 35 in summer, with 16 species common to both seasons, primarily resident birds (Figure 3A).

**Figure 3:**
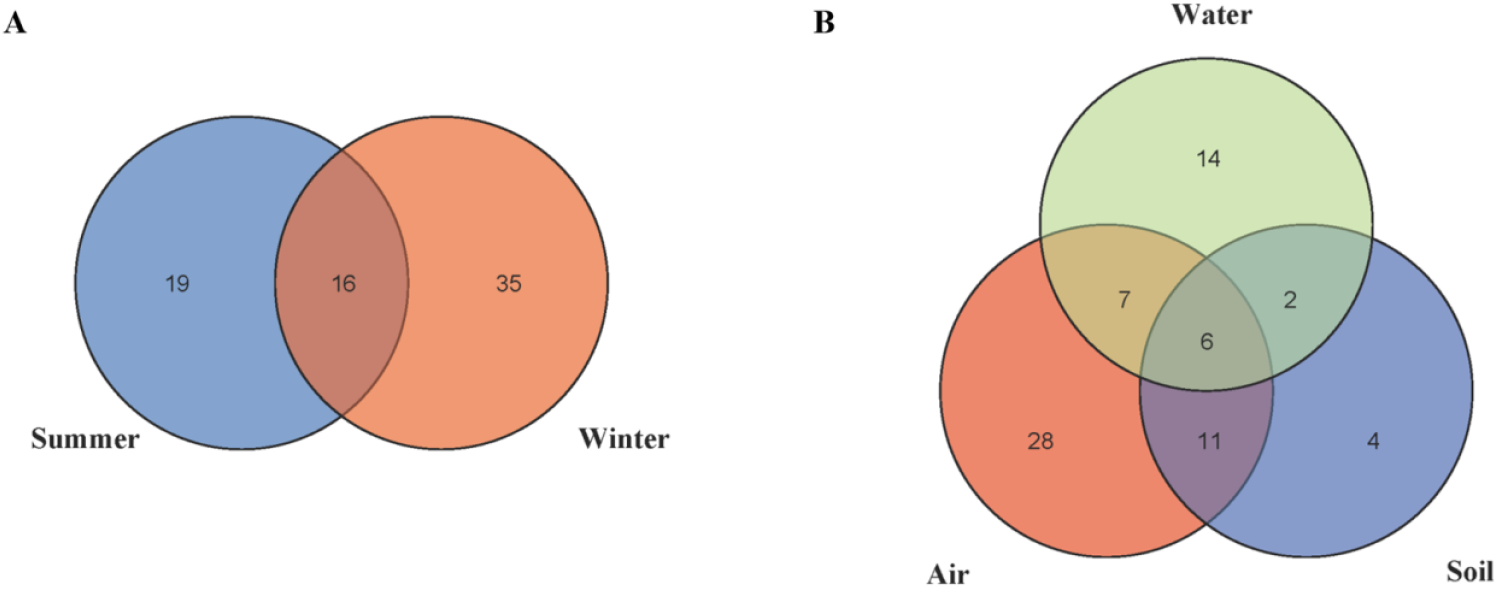
Venn diagram of bird species detected in summer and winter. (A) Venn diagram showing the number of bird OTUs detected in summer and winter; (B) Venn diagram showing the number of bird OTUs detected in water, soil, and air samples.

The positive detection rates varied significantly among the environmental samples (Table 2). Air samples had the highest detection rate, with 40 samples collected across both seasons, of which only 2 samples failed to yield any bird. The average number of species detected per air sample was also the highest. Soil sample results showed seasonal variation, with an 85% positive detection rate in summer, contrasting sharply with a 50% rate in winter. However, it remains unclear whether this discrepancy is seasonal or due to random factors. Despite differing positive rates, the number of bird species detected in both seasons was similar, with 15 species each, although the specific species composition varied.

**Table 2:** Detection of bird OTUs in different types of environmental samples.

| Season | Sample type | Sample size | Number of positive samples | Total number of OTUs | Number of OTUs in water birds | Number of OTUs in forest birds | Weighted average number of OTUs in the sample |
| --- | --- | --- | --- | --- | --- | --- | --- |
| Summer | Water | 21 | 16 | 4 | 3 | 1 | 1.3 |
|  | Soil | 22 | 18 | 15 | 9 | 6 | 3.4 |
|  | Air | 20 | 18 | 30 | 12 | 18 | 10.1 |
| Winter | Water | 22 | 14 | 29 | 18 | 11 | 6.0 |
|  | Soil | 23 | 11 | 15 | 9 | 6 | 2.6 |
|  | Air | 20 | 20 | 34 | 18 | 16 | 15.0 |

Water samples exhibited distinct seasonal trends. During summer, the detection rate was 75%, but only four species were identified (Anas, Egretta, and Columba genera). In winter, the detection rate dropped to 63%, yet 29 species were recorded, predominantly migratory water birds, aligning with expected patterns. These findings suggest a clear seasonal variation in water sample results, with winter displaying significantly greater bird diversity.

Overall, air samples emerged as the most effective medium for investigating bird diversity using eDNA technology, as birds are the only major animal group capable of flight. While water samples have traditionally been favored for biodiversity investigation in aquatic environments, air samples should take precedence when studying birds. Nonetheless, water samples remain vital for detecting waterfowl that prefer wetland habitats. Optimal sampling times based on migratory patterns should be considered, and simultaneous collection of both air and water samples would enhance the detection of major bird species. Given the limited contribution of soil samples to bird detection, their collection may be minimized or focused on areas of high avian activity for cost-effectiveness.

The significant differences in detection frequencies among various bird species were apparent. Among forest birds, the light-vented bulbul and Swinhoe’s white-eye were most frequently detected, while the Chinese pond heron, grey heron, great egret, and black-crowned night heron topped the list for water birds. These common resident species are abundant and easily detectable. Conversely, rarer species, such as the black-faced spoonbill, striated heron, and cinnamon bittern, were detected in limited numbers. Given that samples were collected only once per season, the results reflect inherent randomness, underscoring the high sensitivity of eDNA technology. To increase the detection of rare species, enhancing sampling frequency and reducing randomness is crucial, a challenge shared with traditional survey methods.

Regarding the accuracy of species identification, all species recognized at the species level matched those observed in traditional surveys within the reserve. However, the overall number of bird species detected through eDNA technology was significantly lower than historical records for the reserve, with some common species missing. Possible explanations include: 1) A lack of eDNA primers specifically designed for bird surveys; the universal primers used in this study may not sufficiently cover all taxa; and 2) The non-fixed movements of birds, particularly migratory species, result in some only making brief stops, leading to missed detections due to insufficient sampling frequency.

### 3.2 Analysis of Birds in Biological Samples

The biological samples, primarily feces and feathers, provided more direct evidence of bird presence compared to environmental samples. These samples contained significantly higher nucleic acid levels, facilitating straightforward species identification. In total, 13 bird species were detected from 18 fecal samples, while 8 species were identified from 13 feather samples. This suggests that biological samples can also adequately reflect local bird diversity.

Comparative analysis indicated that all bird species identified in biological samples were also detected in environmental samples, demonstrating the feasibility of using environmental samples for bird diversity assessments. Each fecal and feather sample was individually tested, but the study also explored mixing multiple samples prior to DNA extraction. The resulting bird species composition aligned with expectations, confirming that mixed sampling can be a cost-effective survey method. Depending on practical considerations, pooling several to a dozen fecal or feather samples could effectively represent local bird species composition. However, due to limitations in sample quantity, the study did not determine the optimal number of samples that could be mixed or the ideal mixing quantity.

## Notes

### Competing Interest Statement

The authors have declared no competing interest.

